# Unlocking the Secrets of the Primate Visual Cortex: A CNN-Based Approach Traces the Origins of Major Organizational Principles to Retinal Sampling

**DOI:** 10.1101/2023.04.25.538251

**Authors:** Danny da Costa, Lukas Kornemann, Rainer Goebel, Mario Senden

**Author notes:** Corresponding author(s). E-mail(s); Contributing authors.

## Abstract

Primate visual cortex exhibits key organizational principles: Cortical magnification, eccentricity-dependent receptive field size and spatial frequency tuning as well as radial bias. We provide compelling evidence that these principles arise from the interplay of the non-uniform distribution of retinal ganglion cells (RGCs), and a quasi-uniform convergence rate from the retina to the cortex. We show that convolutional neural networks (CNNs) outfitted with a retinal sampling layer, which resamples images according to retinal ganglion cell density, develop these organizational principles. Surprisingly, our results indicate that radial bias is spatial-frequency dependent and only manifests for high spatial frequencies. For low spatial frequencies, the bias shifts towards orthogonal orientations. These findings introduce a novel hypothesis about the origin of radial bias. Quasi-uniform convergence limits the range of spatial frequencies (in retinal space) that can be resolved, while retinal sampling determines the spatial frequency content throughout the retina.

## 1 Introduction

Primate visual cortex is famously characterized by retinotopic organization; a topology-preserving mapping of visual space onto the cortical surface, such that neurons whose receptive fields are adjacent in visual space are themselves adjacent in the cortex [1–4]. Interestingly, retinotopy forms the basis for a number of other prominent organizational principles of the visual cortex: cortical magnification, receptive field size and spatial frequency tuning are all directly related to eccentricity [1, 2, 5–8]. Specifically, a large proportion of neurons with small receptive fields and preference for high spatial frequencies is dedicated towards processing central vision, whereas progressively fewer neurons with large receptive fields and preference for low spatial frequencies are dedicated towards processing peripheral vision. Additionally, orientation tuning in early visual cortex is directly related to polar angle, with a preference for radial over orthogonal orientations [9–12]. While these organizational principles are well-established, their origin remains insufficiently understood.

It has been hypothesised that cortical magnification is primarily explained by the decrease in retinal ganglion cell (RGC) density, with increasing eccentricity [13–16]. The distribution of retinal ganglion cells exhibits a densely distributed central region, with an incrementally decreasing density, as eccentricity increases [14, 17, 18]. This results in dense sampling of the central visual field, sparse sampling of the periphery, and radial stretching (barrel distortion) of visual space [2, 19]. We suggest that the non-uniform sampling of visual space in conjunction with an almost constant convergence rate from retinal ganglion cells (RGCs) to the cortex[14, 20] underlies not only cortical magnification but all of the aforementioned retinotopy-related organizational principles (Fig. 1). Consider two receptive fields, located at different eccentricities, both comprised of the same fixed number of retinal ganglion cells (RGCs). The low-eccentricity receptive field will cover a small portion of the visual field, as it is constructed from densely packed RGCs. Conversely, the high-eccentricity receptive field will cover a large portion of the visual field, as it is constructed from sparsely packed RGCs. This simple mechanism accounts for increases in receptive field size as a function of eccentricity (Fig. 1b). Furthermore, the increased preference for low spatial frequencies, as eccentricity increases, may be explained by the fact that high frequencies are increasingly compromised by reducing RGC densities (low-resolution sampling of visual space; Fig. 1b). Similarly, consider two receptive fields with a 45°orientation tuning that are respectively located at polar angles of 45°and 135°of visual angle, and how a barrel distortion affects a 45°edge that falls within each of these receptive fields. The edge is radially aligned with the first receptive field and, hence, merely stretched. However, the edge is oriented orthogonally relative to the second receptive field and will thus be bent outward. Since receptive fields in early visual cortex are generally assumed to be Gabor-like filters [21–25], which respond maximally to straight lines, the same edge will elicit a stronger response from the first receptive field compared to the second (Fig. 1c). Radial bias [9, 10, 12] may thus also be caused by RGC sampling.

**Fig. 1.**
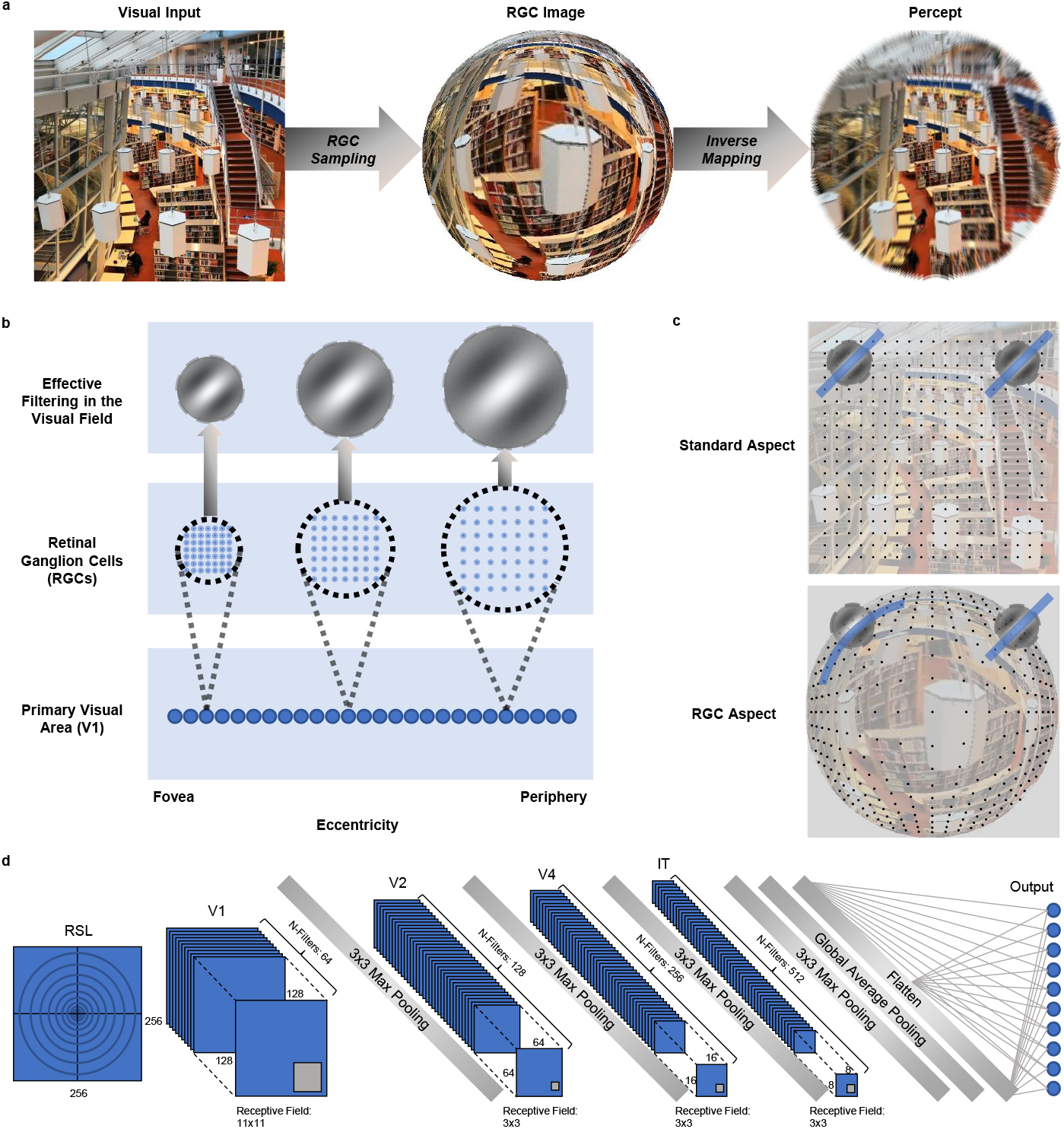
Retinal ganglion cell (RGC) sampling. **a**, Retinal sampling of a square image. The input image (left) is sampled according to RGC distributions, resulting in an RGC representation that is forwarded to the first convolutional layer. The image shows an enhanced central region (foveal vision) with progressive loss of information as eccentricity increases (peripheral vision). Inverting the sampling process transforms an RGC image into a representation similar to human perception. **b**, Schematic overview, demonstrating how an eccentricity-related decrease in RGC density produces progressively larger receptive fields. This results in a progressively lower spatial frequency tuning for an identical V1 filter. **c**, Schematic representation of the radial bias hypothesis. The visual field distortion induced by the RGC sampling induces curved non-radial lines, leading to sub-optimal filter activation for stimuli oriented orthogonal to the radial line. **d**, Schematic overview of the RSL-CORnet-Z model.

We utilize convolutional neural networks (CNNs) to test whether eccentricity-dependent RGC density, combined with a location-invariant convergence rate from the retina to the cortex [as suggested by 14] can give rise to eccentricity- and polar-angle-dependent organizational principles of primate early visual cortex. Due to weight sharing, the receptive field size in CNNs (filters) is location-invariant and thus optimally suited for our purposes. We introduce a retinal sampling layer (RSL) which resamples natural images according to RGC density (Fig. 1a) before they are further processed by the biologically inspired CORnet-Z[26] architecture (Fig. 1d). We show that introducing the RSL leads the CORnet-Z model to develop all of the hypothesized organizational principles. These results simultaneously lend credence to the conjecture that these characteristics emerge in primate visual systems due to retinal ganglion cell distributions and render CNNs more biologically plausible. In addition, our results predict that radial bias is not a general phenomenon, but varies with spatial frequency. Specifically, our results indicate that radial bias is only evident in stimuli with high spatial frequency, while low spatial frequency stimuli give rise to an orthogonal bias.

## 2 Results

### 2.1 Model Training and Evaluation

We chose CORnet-Z [26] as our CNN architecture because it is a biologically inspired CNN with high feedforward simplicity whose four convolutional layers correspond to human V1, V2, V4, and IT. We expanded CORnet-Z by introducing a retina layer (RSL) as the first layer of the model. The resulting RSL-CORnet-Z architecture (Fig. 1d) was trained on upscaled and zero-padded natural images (2048-by-2048-by-3) from ten randomly chosen ImageNet2014 classes[27]. The visual field coverage of these images was set to 20°of visual angle (see Methods for details and Fig. 1a for an illustration). After successful training of the architecture and achieving 88.90% test-set accuracy, we investigated its organizational principles.

### 2.2 Cortical Magnification and Eccentricity-Dependent Receptive Field Sizes

We first asked whether our architecture exhibits cortical magnification and eccentricity-dependent receptive field sizes. To answer these questions, we adopted a population receptive field (pRF) mapping procedure [6] typically used in neuroimaging studies. We exposed the network to oriented (0°, 45°, 90°, and 135°) bar apertures that reveal gratings of different orientations and spatial frequencies. These gratings drive unit responses akin to how flickering checkerboards drive neural responses. Since we were only interested in location-but not feature-specific responses, we averaged layer-specific unit responses across feature maps to obtain stimulus-related *population maps*. These maps thus represent the average activity across all units in one layer that receive input from the same visual field location. Subsequently, we averaged population maps across all gratings presented for every aperture position to obtain population units’ activity in response to each aperture position. The distribution of responses of a population unit over bar apertures constitutes that population unit’s response profile. We compared each population unit’s response profile to the response profiles of a set of candidate Gaussian receptive fields, exposed to the same bar apertures. We consider the Gaussian whose response profile correlates best with the response profile of a population unit as that unit’s receptive field (Fig. 2a).

**Fig. 2.**
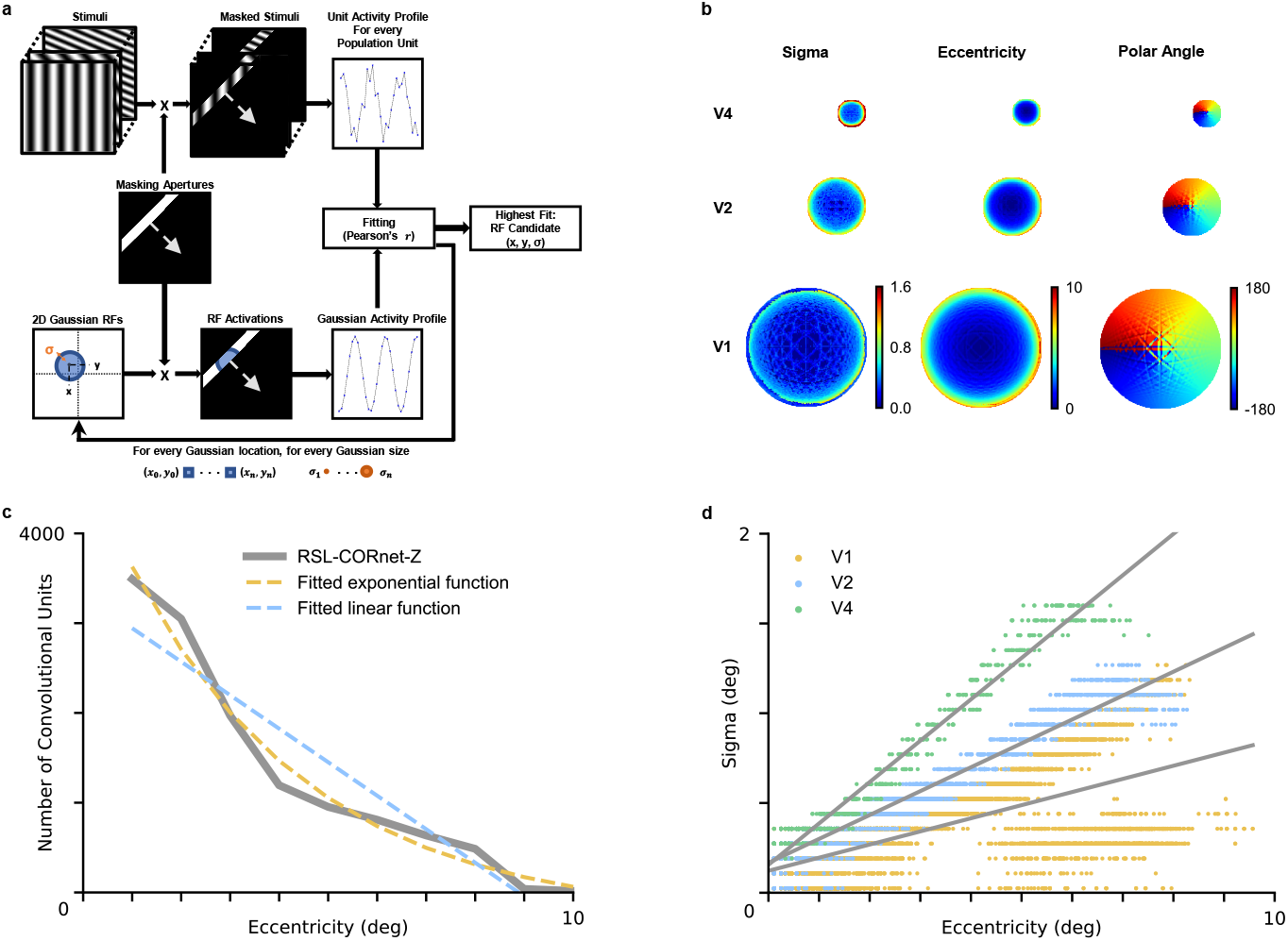
Population receptive field mapping. **a**, Schematic overview of the adapted pRF mapping method used for neural networks.**b**, Retinotopic maps of convolutional layers. Each pixel in the maps reflects a population unit in the corresponding convolutional layer. **c**, Convolutional units per level of eccentricity for the V1 layer, including the best fitting linear and exponential decay functions. **d**, Receptive field size as a function of eccentricity for V1, V2, and V4 layers. Included in gray are the best linear fits for each layer.

By adapting the pRF method to CNNs, we obtained retinotopic maps for CORnet-Z (Fig. 2c). We used the eccentricity data to investigate cortical magnification for the V1 layer by determining the amount of population units per level of eccentricity. To that end, we divided the eccentricity map into contiguous bins (each 1°in width) and counted the number of population units per bin. As expected, density (D) was high at low eccentricities and dropped off as eccentricity (E) increased. We fitted a linear (*D* = 3917.77*e*^(−0.27*∗E*)^− 289.777 *R*^2^ = .88, *F* (10) = 60.52, *p <* .001) and an exponential decay function (*D* = 3917.77*e*^−0.27*E*^ − 289.78; *R*^2^ = .98, *F* (10) = 151.37, *p <* .001) to the number of cells per eccentricity bin (Fig. 2b). The exponential decay function shows a better fit in terms of explained variance(*R*^2^ = .98). These observations qualitatively agree with cortical magnification in primate vision[1, 28, 29].

We also used retinotopic maps to investigate whether receptive field sizes increase as a function of eccentricity. To this end, we utilized the slope and intercept of the fitted line that best captures the relation between pRF receptive field sizes and eccentricity (Fig. 2d).In the V1 layer, receptive field sizes split into two distinct curves as eccentricity increases. This is likely due to receptive fields in the visual field becoming increasingly elliptical with eccentricity as a result of the barrel distortion (see Supplementary Fig. 2). Our observations support previous reports of radial elongation of peripheral receptive fields in monkeys and cats [30, 31]. In addition, it has previously been shown that pRF mapping with circular Gaussians produces a mix of over- and under-estimated receptive field sizes when receptive fields are elliptical [32] as it captures either their semi-minor or semi-major axis. Given that over- and under-estimates are nearly balanced and that the geometric mean of the sizes of semi-minor and semi-major axes approximates the size of a circular receptive field, we nevertheless fit a single line through the data *RFsize_V_* _1_ = 0.073*E* + 0.122 (*R*^2^ = .47, *F* (12644) = 11319.34, *p <* .001). The eccentricity-related V1 receptive field sizes for macaque monkeys, as reported by Freeman and Simoncelli [33], exhibit a shallower slope and a higher intercept (*RFsize_Monkey__V_* _1_ = 0.042*E* + 0.271). This higher intercept suggests that the filter size of the convolutional units is larger compared to biological receptive fields. Remarkably, the size-eccentricity slope of the RSL-CORnet-Z V1 layer (0.073) is highly similar to the slope of 0.072 originally described by Hubel and Wiesel in monkey primary visual area V1 (*RFsize* = 0.072 *· RFeccentricity* + 0.017)[34, 35].

We also performed linear fits to the pRF data of the V2 and V4 layers. The results indicate a high fit in terms of explained variance for both V2 (*RFsize_V_* _2_ = 0.133*E* + 0.167, *R*^2^ = .92, *F* (3094) = 38110.84, *p <* .001) and V4 (*RFsize_V_* _4_ = 0.230*E* + 0.156, *R*^2^ = .96, *F* (726) = 19575.874, *p <* .001) layers. Comparing these model slopes to monkey V2 (0.119E) and V4 (0.199E) slopes [33], it becomes apparent that while the exact slope values do not match due to the size of our convolutional filter, our model exhibits a similar progression in slopes from V1 to V4 as the monkey. Specifically, the V4 slope is larger than the V2 slope by a factor of 1.73 in our model and by a factor of 1.67 in the monkey. It is important to note that the effective receptive field size of the convolutional filter in the visual field depends on several factors, including the resolution of the output image of the RSL, the FOV that is modeled by the RSL, and the receptive field size of the convolutional layer. It is therefore principally possible to optimize these parameters to match the receptive field size of the model arbitrarily well to biological receptive field sizes by manipulating one, or several, of these parameters. As such, the quantitative fit of the size-eccentricity relationship in our model with empirical data should not be over-interpreted. Instead, we wish to emphasize the qualitative observation that receptive field sizes increase as a function of eccentricity.

### 2.3 Eccentricity-Dependent Spatial Frequency Tuning

Next, we explored the possibility that the RGC distributions can account for eccentricity-dependent spatial frequency tuning. We investigated whether such a relationship emerges in the network by presenting sinring apertures [cf. 36, 37] of various spatial frequency-eccentricity combinations to the RSL-CORnet-Z (Fig. 3a). Here, the spatial frequencies are expressed as cycles per radian (cpr). As hypothesized, the spatial frequency preference of population units shifted with eccentricity. More precisely, for units encoding central regions of the visual field, the highest spatial frequency 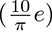 elicited the strongest response. For population units encoding peripheral regions of the visual field, the lowest spatial frequency 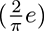 elicited the strongest response (Fig. 3b).

**Fig. 3.**
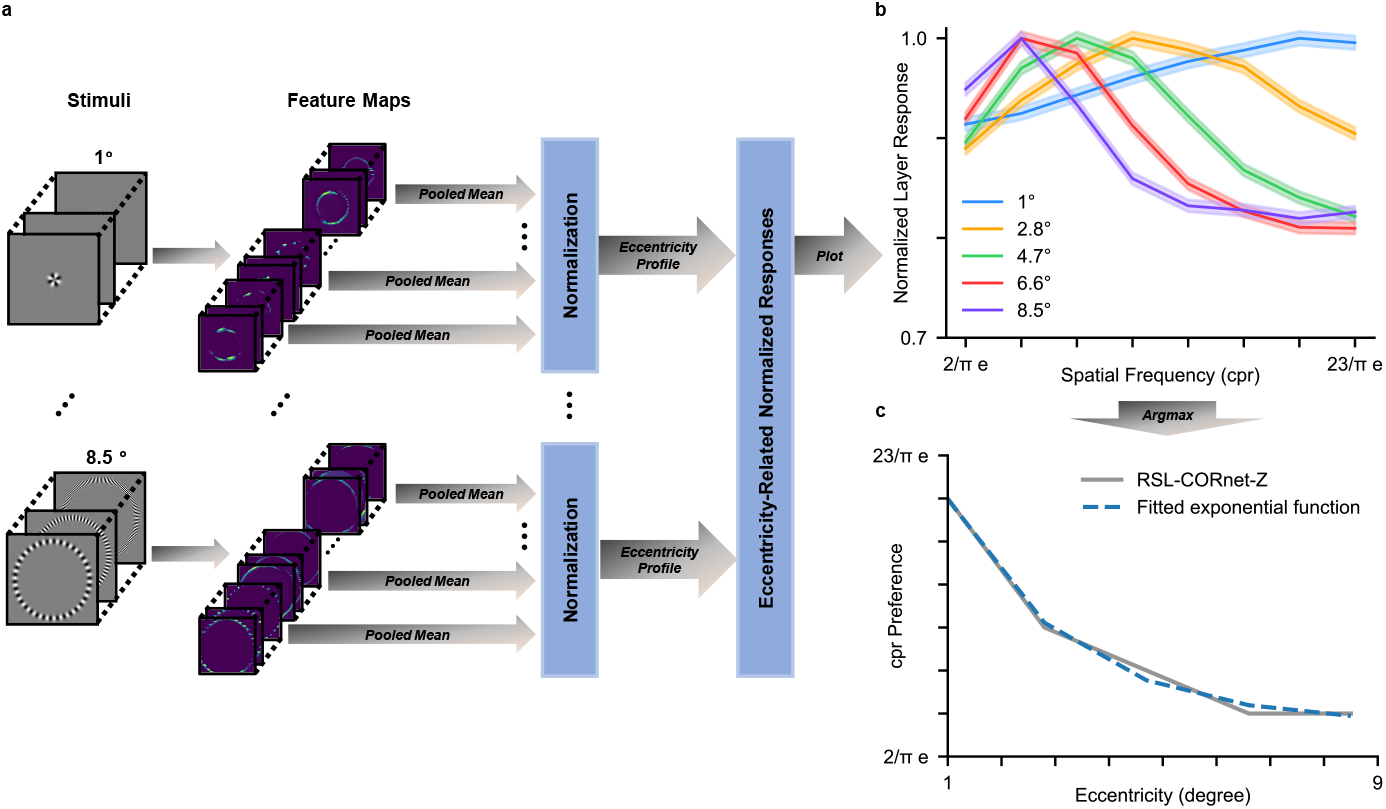
Eccentricity dependent spatial frequency tuning. **a**, Schematic overview of the Sinring method for detecting spatial frequency tuning at different levels of eccentricity in neural networks. **b**, Normalized averaged layer response for the V1 layer as a function of spatial frequency. Responses observed at five eccentricities are presented separately (color coded). The shaded area surrounding each curve represents the 95% confidence interval. **c**, Spatial frequency preference as a function of eccentricity.

Consistent with previous findings in primate visual systems [5, 7, 36], the RSL-CORnet-Z displays an exponential decay relationship between eccentricity and spatial frequency preference in the V1 layer (*cpr_pref_*= 48.82*∗e^−^*^0.44*x*^ + 8.54, *R*^2^ = .99, *F* (5) = 161.98, *p* = .006), as illustrated in Figure 3c. The change in spatial frequency preference for fixed CNN filters is a direct consequence of the non-uniform retinal sampling of visual input. In the fovea, high-spatial frequencies are expanded into the preferred frequency range of the filter, whereas in the periphery, increasingly low-spatial frequencies are compressed into the preferred frequency range of the filter. Thus, a filter’s fixed spatial frequency tuning in retinal or cortical space results in a decrease in spatial frequency tuning as a function of eccentricity in the visual field. Our findings are consistent with findings that primate V1 cells exhibit constant spatial frequency tuning in cortical space, regardless of visual field eccentricity [36, 38].

### 2.4 Radial Bias

Lastly, we investigated the presence of radial bias in the V1 layer of the network by presenting full-field sinusoidal gratings of eight orientations at a spatial frequency of 4 c/deg [cf. 9]. We created different stimuli per level of orientation by varying the phase offset of the grating. As before, we averaged across feature maps to obtain one population map. Subsequently, we averaged all population maps per grating orientation, resulting in *orientation maps* (Fig. 4a). Subsequently, units in the orientation maps were divided into 16 45° polar angle bins. (Fig. 4b). We computed the mean activity for each of these bins (Fig. 4c). In line with radial bias observed in the primate visual cortex, the semi-major axis of the resulting ellipse aligns with the orientation of sinusoidal gratings.

**Fig. 4.**
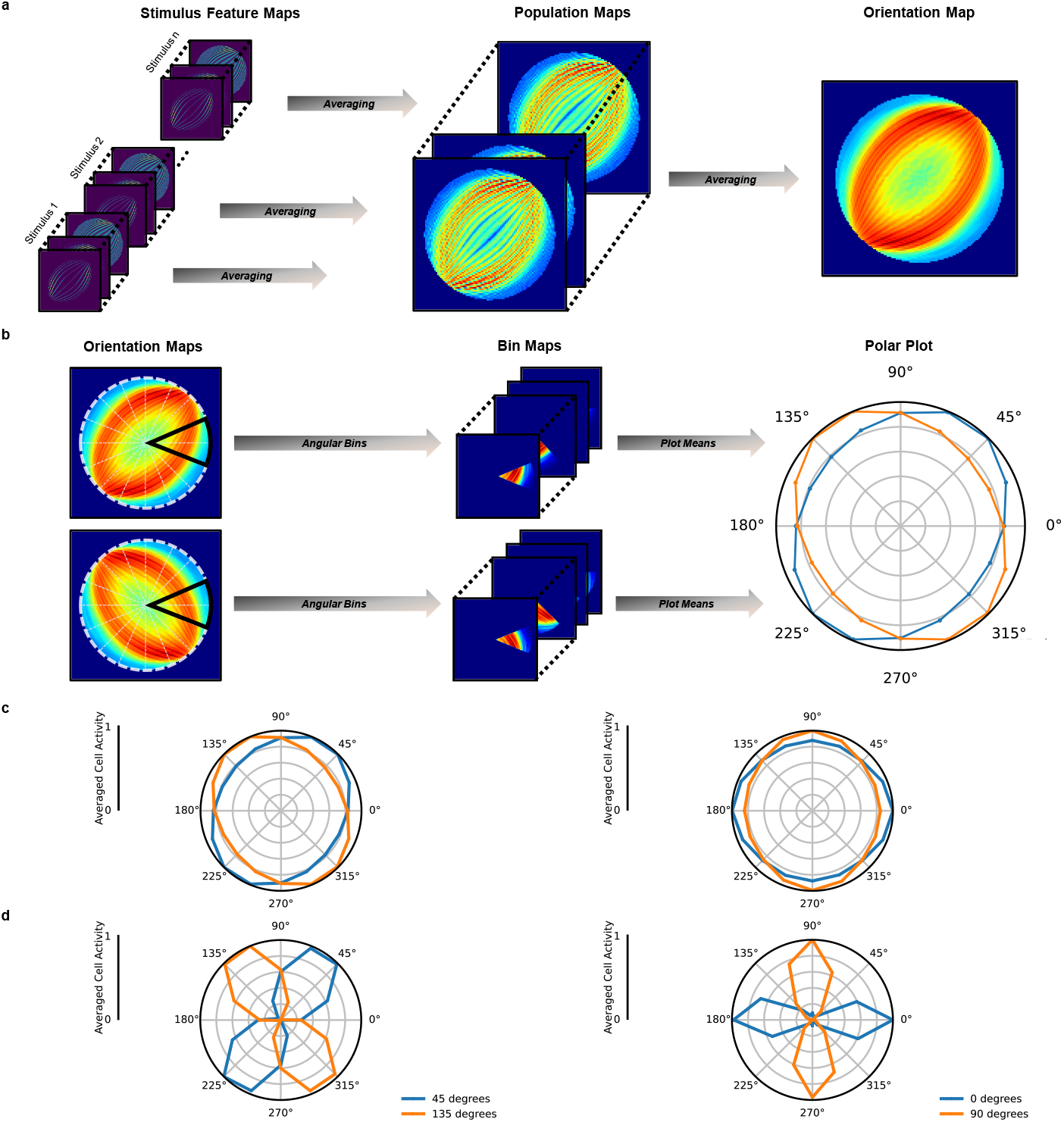
Radial bias. **a**, Feature maps for each stimulus are averaged into population maps. The mean feature maps of stimuli belonging to the same stimulus orientation are averaged into a single orientation map. **b**, Resulting maps are then split into 16 overlapping polar angle bins with each bin covering 45°. Total activity within each bin is then max-normalized and displayed in a polar plot. **c**, Polar plots displaying radial bias for 4 c/deg sinusoidal gratings. The left plot shows the result for 45° vs 135° sinusoidal gratings, the right plot shows 0° vs 90°. The data is normalized per stimulus orientation. **d**, Same data as in c, with min-max normalization per stimulus orientation to visually highlight the preferred orientation. Note that this normalization procedures exaggerates the magnitude of the bias

We quantified the existence and direction of an orientation bias (i.e., a preference for one orientation) in terms of the ratio of the total unit activity (*A*) along the axis aligned with the grating orientation to the total unit activity along the axis orthogonal to the grating orientation. Specifically, we computed *A_grating_/A_orthogonal_*, where *A_grating_*is the sum of averaged unit activities in the bins with a 0°and 180°angle to the grating orientation. Similarly, *A_orthogonal_* is the sum of average unit activities in the bins with a 90°degree and 270°angle to the grating orientation. A ratio close to unity indicates the absence of any orientation bias. Ratios larger than unity indicate an increasingly strong radial bias, whereas ratios smaller than unity indicate an increasingly strong orthogonal bias. Our analysis revealed the radial bias ratios for the various grating orientation. At a spatial frequency of 4 c/deg, the radial bias ratio was 1.16 at a grating orientation of 45°, 1.24 at 135°, 1.14 at 0°, and 1.14 at 90°(Fig. 4c). The average radial bias ratio of 1.17 for gratings at a spatial frequency of 4 c/deg demonstrates a general radial bias in the V1 layer.

### 2.5 Radial Bias Depends on Spatial Frequency

Radial bias experiments typically use high spatial frequency gratings (*≥* 3 cycles per degree; c/deg) [c.f. 9, 11]. This implicitly assumes that radial bias is independent of the spatial frequency content of the input. To test whether this assumption holds in the network, we repeated our experiment with varying spatial frequencies of the sinusoidal gratings (0.25, 0.5, 1, 2, 3, 4, 5 c/deg). Results for 0.25, 2, and 5 c/deg, for 0°and 90°gratings are presented in Fig. 5a. Results for 45°and 135°gratings are presented in Fig. 5b. Results for all other spatial frequencies and grating orientations can be found in Supplementary Fig. 3 and Supplementary Fig. 4.

**Fig. 5.**
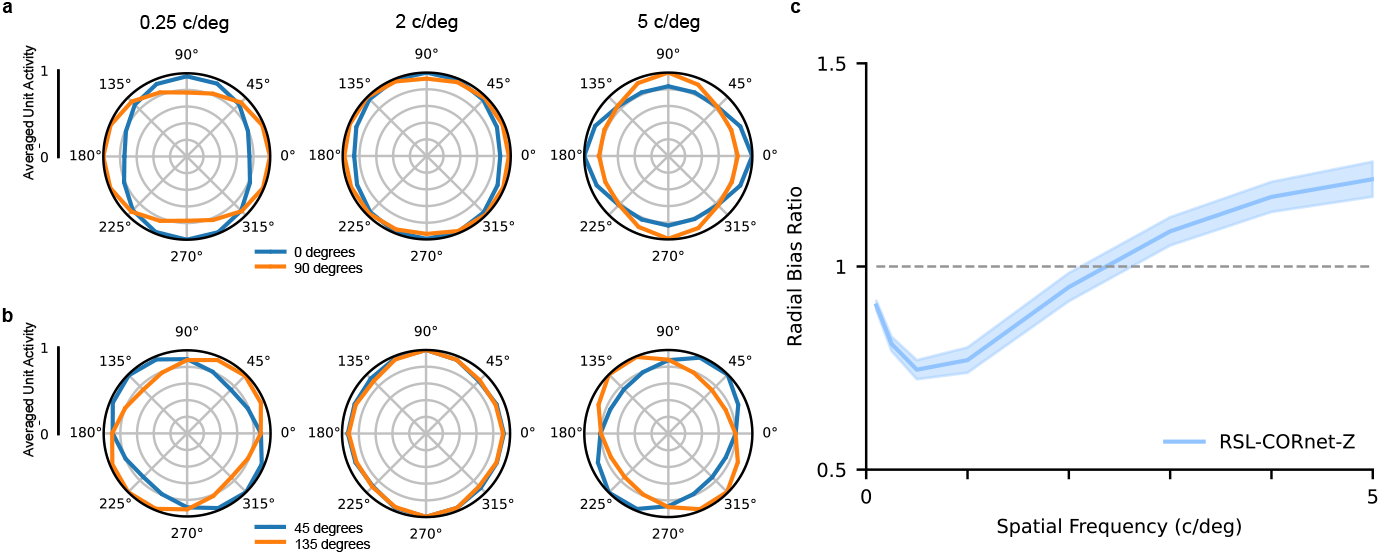
RSL-CORnet-Z orientation biases. **a**, Orientation biases for 45 and 135° sinusoidal gratings for 0.25, 2, and 5 cycles per degree (c/deg). **b**, Orientation biases for 0° and 90° stimuli for 5, 20, and 40 c/deg. **c**, Averaged radial bias ratio per level of spatial frequency (c/deg). A value of 1.0 indicates the absence of any orientation bias. A value below 1.0 indicates a orientation preference for orthogonal orientations, while a value above 1 indicates the presence of radial bias. The shaded area displays the 95% confidence interval.

To determine the radial bias as a function of spatial frequency, we computed the mean radial bias ratios for every level of spatial frequency. Surprisingly, our results reveal that radial bias in the V1 layer depends on the spatial frequency of the sinusoidal gratings. While a broad range of mid-to-high spatial frequency gratings gives rise to radial bias, this bias disappears, and an orthogonal-to-radial bias emerges once the spatial frequency of the gratings is sufficiently reduced (Fig. 5c). In other words, for low spatial frequency stimuli, units in the V1 layer of the RSL-CORnet-Z exhibit a preference for orthogonal over radially oriented lines.

### 2.6 Orientation Bias Exploration

To gain a deeper insight into the mechanisms behind radial bias, as well as its reversal for low spatial-frequency gratings, we investigated our original hypothesis which stated that radial bias emerges due to curving of non-radial lines. We hypothesized that V1 (in the primate and our network) exhibits Gabor-like filters that show a higher response to straight lines compared to curved lines. Given that sampling by the RSL curves edges that are not aligned with the radial axes, Gabor-like filters should demonstrate reduced responses for non-radial stimuli, thus accounting for radial bias. To investigate this hypothesis, we created seven-by-seven-by-three pixel images with a single radial line of various orientations (0°, 45°, 90°, and 135°) and parametrically varied the curvature of these lines in both orthogonal-to-radial directions. In addition, we created 3D Gabor kernels matched to the spatial frequency of our artificial stimuli, and with orientations from 0° to 180° degrees, in 16 steps. We measured the cosine similarity between the Gabor kernels and the curvature stimuli, per level of line curvature. For comparison, we also measured cosine similarity between RSL-CORnet-Z filters and the curvature stimuli. As can be appreciated from Fig. 6a, the Gabor filters show a clear drop in cosine similarity when lines get more curved. This confirms our conjecture that typical Gabor filters have an impaired performance when dealing with curved stimuli. Surprisingly, however, the model filters fail to show this effect. If anything, they reveal a small trend in the opposite direction. This indicates that the CNN had sufficient degrees of freedom to compensate for the presence of curved edges by developing curved filters (see Supplementary Fig. 5). The observation that the CNN nevertheless exhibits radial bias points to an entirely different mechanism.

**Fig. 6.**
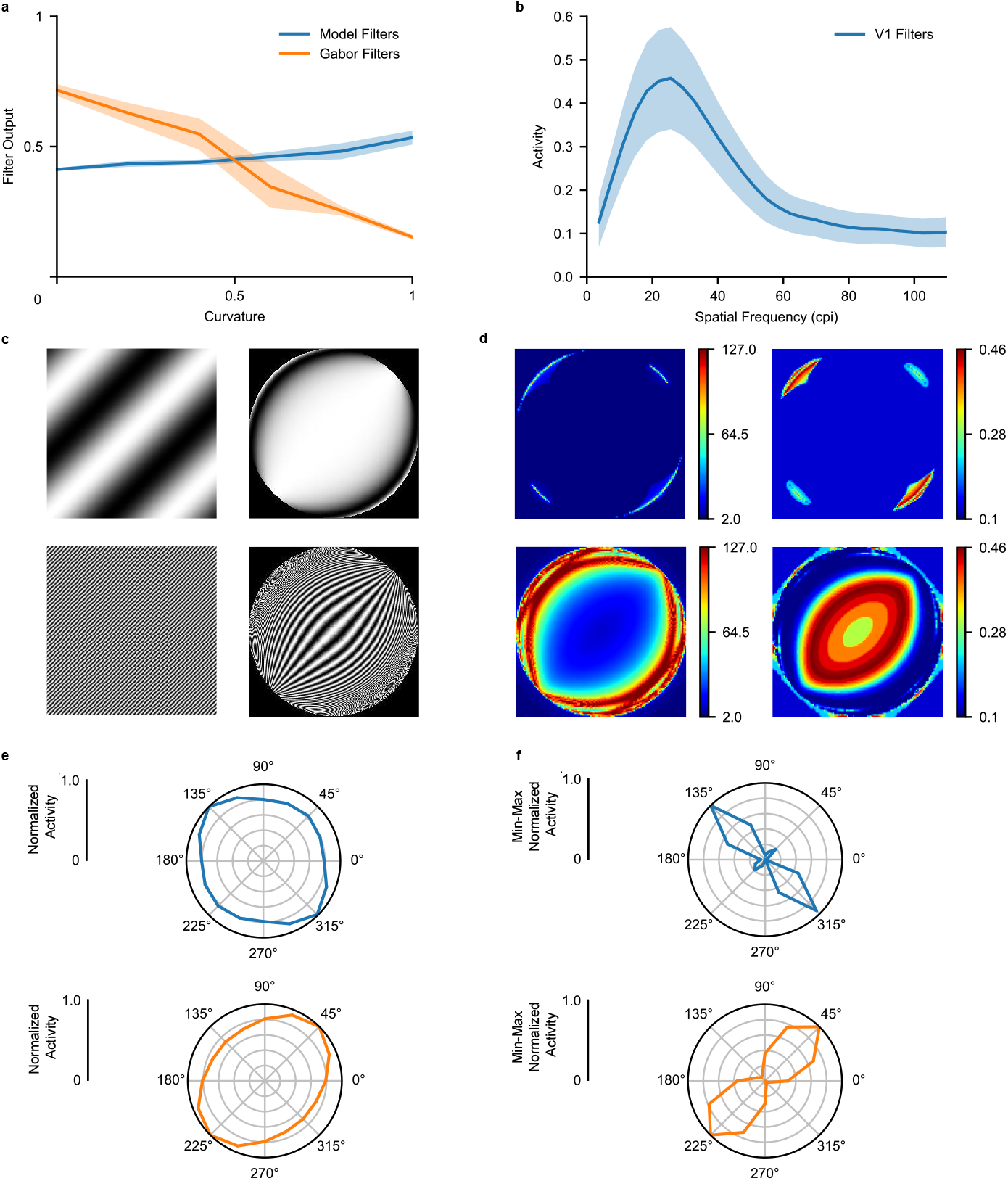
Orientation bias analysis. **a**, V1 filter, and Gabor responses to curved lines. A curvature value of zero indicates a straight line, one indicates maximum curvature. **b**, Average response of V1 filters to gratings (immediately placed within the resampled image) varying in cycles per image (cpi), orientation and phase. Responses are averaged per level of spatial frequency (cpi). **c**, The left column displays the original stimuli used for spatial frequency analysis, the right column displays the same stimuli after ganglion sampling. Low and high spatial frequency stimuli are displayed in the top and bottom row, respectively. **d**, The left column displays the local spatial frequency content (extracted with a Gabor filter pyramid) of the stimuli displayed in the left column of c. The right column shows the estimated V1 layer response, given the local spatial frequency content. **e**, Normalized orientation bias based on the right column of d. **f**, Identical to e, but min-max normalized.

Further investigations revealed that an orientation bias, whether radial or orthogonal, emerges from the combination of limited spatial frequency tuning in the retinal space and the effects of resampling on the spatial frequencies across the visual scene. The limited spatial-frequency tuning in retinal space results from the retina’s fixed convergence rate to a V1 cell. This means that only specific spatial frequencies can be resolved in the resampled image (Fig. 6b). The spatial frequencies present in the resampled image depend on the spatial frequency content of the original stimulus. Simply put, low spatial frequencies are compressed into the appropriate frequency range at different locations than high spatial frequencies. To show this, we analyzed gratings’ local spatial frequency content in the resampled image space. As can be appreciated from Fig. 6d (top row), low spatial frequencies in the original image are transformed into the frequency range to which the network is sensitive, mainly within regions that are orthogonal to the grating’s orientation. By contrast, high spatial frequencies in the original image are transformed into the appropriate frequency range in an elongated region whose semi-major axis is aligned with the grating’s orientation (Fig. 6d bottom row). Thus, by virtue of their size, network filters can only resolve frequencies within a narrow range, and this frequency range is limited to specific regions of the sampled image. We suggest that, due to quasi-uniform convergence, neurons in V1 likewise have a limited range of frequencies they can resolve in retinal space. Depending on the original stimulus, frequencies are differentially distributed across the retina. For high spatial frequency gratings, typically employed in radial bias experiments, these regions happen to align with the orientation of the grating.

## 3 Discussion

This study presents compelling evidence that key organizational principles in the primate visual cortex result from the non-uniform distribution of retinal ganglion cells in conjunction with a quasi-uniform convergence rate from retinal to cortical neurons. The non-uniform distribution of retinal ganglion cells causes a barrel-distortion of visual space such that foveal regions are enlarged, whereas peripheral areas are compressed in retinal space. Quasi-uniform convergence from retinal to cortical neurons implies that all V1 neurons receive input from a constant amount of retinal ganglion cells. Given that most retinal ganglion cells are concentrated near the fovea, quasi-uniform convergence implies that disproportionally many V1 neurons are tuned towards foveal representations, thus accounting for cortical magnification. Furthermore, the spacing between ganglion cells increases with eccentricity. Hence, peripheral V1 neurons sample larger regions of visual space, albeit at a lower resolution, than foveal V1 neurons. This accounts for the eccentricity-dependent enlargement of receptive fields tuned to progressively lower spatial frequencies.

Our results further indicate that the interplay of the same two factors gives rise to radial bias. Specifically, quasi-uniform convergence from retinal to cortical neurons means that all V1 neurons have equally sized receptive fields in retinal space. Given that the size of the receptive field fundamentally limits its spatial frequency tuning, all V1 neurons are likely to have highly similar spatial frequency preferences in retinal space, regardless of their location. While quasi-uniform convergence leads to location-invariance of spatial frequency tuning in retinal space, non-uniform sampling of visual space ensures that spatial frequencies themselves are not uniformly distributed across the retina. For a simple grating stimulus, this implies that spatial frequencies in retinal space can only be resolved at specific retinal locations with the exact locations depending on the spatial frequency of the grating. Interestingly, for mid-to-high spatial frequency gratings, these locations form an elliptical region whose semi-major axis is aligned with the orientation of the grating. This manifests as radial bias in our in-silico experiments. For low spatial frequency gratings, however, these locations form two separate regions along an axis that is orthogonal to the orientation of the grating. This manifests as an orthogonal bias in our in-silico experiments. Based on these insights, we make the novel prediction that radial bias in the primate visual cortex is likewise spatial-frequency dependent.

Deep learning offers powerful new opportunities to model biological brain regions and functions. Its utility in this regard arises from its as-of-yet unique integration of brain-inspired computational principles [39–42] with hitherto unseen effectiveness at solving perception tasks [41, 43–45]. This renders deep neural networks well suited to test hypotheses in silico by training them on ecologically relevant tasks and exposing them to stimuli used in neuroscientific experimentation. We utilized CNNs in particular to test the hypotheses that several organizational principles of the early visual cortex arise from the interplay of only two simple factors; non-uniform sampling of visual space across the retina and a quasi-uniform convergence rate from the retina to the cortex. In the present study, non-uniform sampling was assumed to arise from the distribution of retinal ganglion cells. This is an over-simplification given that the geometry of the eye, photoreceptor distributions, the lateral spread of information in the retina via bipolar and amacrine cells as well as the convergence rates from photoreceptors to ganglion cells, from the retina to the lateral geniculate nucleus (LGN), and from the LGN to V1, all contribute to the exact sampling of visual space. By omitting these details, the organizational principles arising in our model may not quantitatively match their biological counterpart. However, even after considering these factors, the sampling of visual space would still result in a barrel distortion. This type of distortion ultimately leads to the organizational principles investigated here such that our results nevertheless provide a qualitative account of the mechanisms that give rise to these organizational principles in the primate visual cortex.

Another limitation of our approach is the weight-sharing inherent in CNNs. In contrast to the primate visual cortex, where locally specialized filters may have arisen through evolutionary processes and visual experience, CNNs apply the same filters everywhere. This is particularly relevant in the context of local curvature arising from retinal sampling. While CNNs apply the same filters everywhere, they may include sufficiently many filters that at least one of them may be specifically tuned to the curvature that arises at any spatial location. Indeed, our results show that the CNN is robust to curvature. Furthermore, contrary to our original conjecture (see Introduction), radial bias does not appear to result from curvature but rather from a fundamental limit on spatial frequency tuning in retinal space combined with distributions of spatial frequency content throughout the retina.

This radically new, frequency-based account of radial bias further highlights that deep learning can be a powerful tool in neuroscience to generate new, testable hypotheses. Specifically, we predict that radial bias is frequency-dependent in the primate visual cortex and only exists for moderate-to high-spatial frequency stimuli. For low spatial frequency stimuli, one should instead observe an orthogonal bias. This prediction can be readily tested with fMRI using the same experimental paradigms that have hitherto been employed to establish radial bias [c.f. 9–11]. One would only need to present oriented gratings with much lower spatial resolution than those typically employed in these experiments.

In conclusion, our results suggest that the interplay of two simple principles, namely, the non-uniform distribution of RGCs and a quasi-uniform convergence rate from the retina to the cortex, give rise to cortical magnification, eccentricity-dependent receptive field sizes and their spatial frequency preferences, and radial bias. Our work contributes in several additional ways. First, it demonstrates how deep learning can be used to both test and generate novel neuroscientific hypotheses. Second, we demonstrate how neuroimaging methods can be adapted and used to investigate convolutional neural networks.

## 4 Methods

### 4.1 Retinal Sampling Layer

The RSL takes a uniformly sampled square image as input (non-square images will be zero-padded along the smaller dimension) and outputs a new, non-uniformly sampled square image. Non-uniform sampling in the output image corresponds to the non-uniform placement of ganglion cells in the retina. In essence, the resampling operation converts between visual field coordinates and ganglion cell coordinates. Since the non-uniform placement of ganglion cells in the retina is eccentricity dependent and does not affect polar angles, it is convenient to use polar coordinates for this conversion. Resampling then reduces to the effect of converting visual field radii (*r_vf_*) to ganglion cell radii (*r_gc_*); i.e., the number of ganglion cells in the interval [0, *r_vf_*]. We can obtain the number of ganglion cells by integrating a function relating ganglion cell density to eccentricity (visual field radius). For this, we use an empirically derived [18] ganglion cell density function:

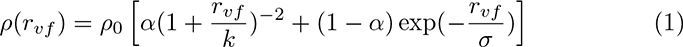

where *ρ*_0_ is the density at *r_vf_*= 0, *k* is the eccentricity at which density is reduced by a factor of four, *σ* is the scale factor of the exponential and *α* is the weighting of the first term. Parameter definition and values are shown in Table 1. The ganglion cell radius corresponding to a given visual field radius is then given by:

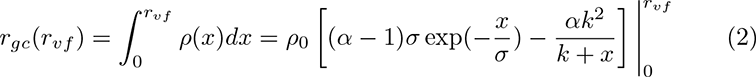

**Table 1.** RGC density function

| Symbol | Value | Definition |
| --- | --- | --- |
| $\rho_0$ | 33162 | ganglion cell density at $r_{vf} = 0$ |
| $r_{vf}$ | 20° | visual field radius in degrees |
| $k$ | 1.05 | $r_{vf}$ at which density is reduced by a factor of four |
| $\sigma$ | 22 | scaling factor in the exponential term |
| $\alpha$ | 1 | weighting of the quadratic term |

This relation allows one to move a pixel in the input image to its corresponding location in the output image. Specifically, after Cartesian pixel coordinates in the square input image are (1) centered at zero by subtracting half of the image width from both the x- and y-coordinate and (2) converted to Polar coordinates, the visual field radius of a pixel is given by 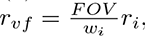 where *FOV* is the visual field of view (degrees) covered by the input image, *w_i_* is the input image width and *r_i_* is the radius of the pixel in the input image space.

The field of view can be anywhere from 5° up to 100°, depending on the distance from which an image may be viewed. The radius of the pixel in the output image space is then 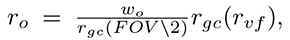 where *w_o_*is the output image width which can be freely chosen. Note that this latter conversion assumes that the convergence rate of retinal ganglion cells to V1 neurons is uniform across the visual field [14].

Such a forward resampling approach may lead to empty pixels in the output image. Therefore, it is preferable to employ an inverse resampling; i.e., to go through all pixels in the output image and query which pixels in the input image they correspond to. This renders it necessary to invert equation 2, which is complicated by the exponential term. However, since the exponential term is mainly necessary to capture ganglion cell density at very large eccentricities, we chose to omit it by setting *α* to 1. Please note that the contribution of the exponential term is generally small since *α >* 0.97 for all results reported by Watson[18]. Therefore, equation 2 becomes:

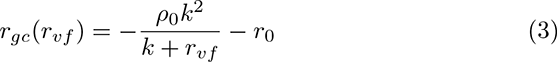

where 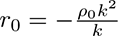 is the number of ganglion cells at zero eccentricity. The visual field radius is then given as a function of the ganglion cell radius

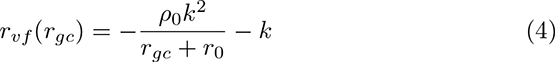

In most cases, pixel coordinates in the input image will not be integers. Therefore, we compute the intensity value of a pixel in the output image as the distance-weighted average over input image pixels whose coordinates are closest to the computed pixel coordinate. For fixed dimensions of input and output images and a fixed FOV, sampling can be performed by summarizing the relationship between pixels in the input and output image in a sparse weight matrix and multiplying this matrix with each of the vectorized RGB channels of the input image.

### 4.2 CORnet-Z model and Natural Images

We used the CORnet-Z convolutional model architecture developed by Kubilius et al.[26] and trained it on natural images taken from 10 randomly chosen classes from the ImageNet2014 training library (n01443537: goldfish, n02132136: brown bear, n03530642: honeycomb, n04004767: printer, n04310018: steam locomotive, n04398044: teapot, n04552348: warplane, n07873807: pizza, n09288635: geyser, n12998815: agaric)[27]. The CORnet-Z model was designed to be a simple abstraction of the human visual system. The model consists of four convolutional ReLU layers, each followed by a max pooling layer. The layers in the model correspond to V1, V2, V4, and IT areas in the human brain. The model’s output consists of an average pooling layer followed by a 10-unit softmax classification layer in which each output node corresponds to one of the ten image classes. Two versions of the model were trained and tested; one CORnet-Z model with the RSL preceding the first convolutional layer(RSL-CORnet-Z) and one standard CORnet-Z model. The hyperparameters for the model’s layers are identical to those described in the original paper[26], except for a change in filter size for the V1 layer from seven-by-seven-by-three to 11-by-11-by-three. Prior to training, weights were initialized by sampling from a truncated normal distribution with a mean of zero and a standard deviation of 0.01. Adadelta with standard values was used for network optimization [46]; the only exception was that the initial learning rate was set to a value of 0.5. Training time was set to a standard 200 training epochs. Training and initialization of hyperparameters were based on a random search method.

Images from the ImageNet2014 library [27] vary in resolution. Therefore, we resized all images such that the largest dimension of each image is equal to 256 pixels. The smaller dimension was zero padded such that the overall resolution of the image conforms to 256-by-256 pixels. In addition to creating a uniform input shape for all images, this method will prevent any stretching or compression of the original image. For the RSL variant of the CORnet-Z model original images were upscaled, using the OpenCV python library [47] and the standard bilinear interpolation, such that the largest dimension of each image is equal to 2048 pixels. We then applied zero padding to the smaller dimension of the image. This resulted in an overall 2048-by-2048 image resolution. The upscaled images were directly fed to the RSL layer and were modeled to represent a FOV of 20°. The RSL output was set to 256-by-256-by-3 pixels. We evaluated the classification accuracy using sparse categorical cross-entropy. We also tested the standard CORnet-Z model for the presence of the aformentioned organizational principles. We present the results in Supplementary Fig. 1. All networks were constructed and tested using Keras in Tensorflow 2.4.0, using NVIDIA CUDA 11.0 with cuDNN v8.2.1 support on a Gigabyte NVIDIA RTX 3080 Aorus Master 10GB graphics card.

### 4.3 Topography Analysis

#### 4.3.1 Retinotopy

We investigated the retinotopic organization of the models by fitting location and size parameters of an isotropic Gaussian population receptive field (pRF) [6] using a grid search method. To that end, we covered the input image with bar apertures for each of four orientations (0°, 45°, 90°, and 135°). The bar was eight pixels wide for 256-by-256 images and had a stride of 4 for each bar presentation. For the upscaled 2048-by-2048 images, the bar width and stride were multiplied by eight. At each orientation-location combination, the bar aperture revealed 1024 sinusoidal gratings. These gratings were created for 8 orientations (0, 22.5, 45, 67.5, 90, 112.5, 135, 157.5 degrees of polar angle) with four different spatial frequencies (0.0735, 0.147, 0.294, 0.5885 cycles per degree; c/deg) and 32 phase offsets (0 to 1.75*π*) per c/deg, per orientation. This resulted in 1024 grating stimuli with 128 examples per grating orientation.

The layer’s location-based visual field activity to each unique bar location was computed in two steps. First, for each bar location, feature maps produced by the convolutional layer were averaged per grating into *population maps*. Every *population unit* in the population map represents the average activity to a single stimulus, for all convolutional units in the layer processing the same visual field location. Second, the population maps for all background gratings per bar location were averaged into an averaged *activation map*. A response profile for every population unit was created by vectorizing the respective unit’s activity for all bar locations.

We constructed candidate receptive fields as isotropic Gaussians at 16,384 uniformly distributed locations with size (*σ*) ranging from 0.025° to 1.6° in 20 linear steps for the grid search. We then obtained a response profile for each receptive field candidate by computing the dot product between the vectorized Gaussian and each vectorized bar aperture. The receptive field for every population unit in the layer was determined by the location and size of the Gaussian whose response profile correlated highest with the response profile of that unit. A schematic overview of the adapted pRF mapping procedure was reported in Fig. 2a.

We utilize the results of the adapted pRF mapping procedure to investigate cortical magnification and the relationship between receptive field size and eccentricity. For cortical magnification, we count the number of population units falling within 1°contiguous bins of visual field eccentricity, discarding population units whose fit (correlation between their response profiles and best candidate receptive field) deviates from the mean fit by three standard deviations or more. Next, we fit a linear and an exponential decay function to the unit counts. To investigate the relationship between receptive field size and eccentricity, we separately fit linear functions to receptive field eccentricity and size estimates of population units in V1, V2, and V4 layers.

#### 4.3.2 Eccentricity-Related Spatial-Frequency Preference

We investigated eccentricity-dependent preferences emerging in the CNN by exposing it to achromatic sine-wave ring stimuli (sinrings) varying in eccentricity and spatial frequency [36, 37]. The rings were presented at five eccentricities (*e*; 1°, 2.8°, 4.7°, 6.6°, and 8.5°; image radius = 10°). To ensure equal spatial frequency content at each eccentricity, the angular spatial frequencies were set to 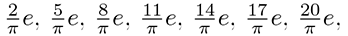 and 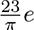 cycles per radian (cpr).

To investigate the response of the network to specific eccentricity-frequency combinations, we first computed population maps by averaging the feature maps of the V1 layer for each sinring stimulus. We then used the pooled mean activity of the population maps from a single stimulus to indicate the layer’s response to that stimulus. Next, we grouped the resulting layer responses based on the level of eccentricity and normalized the data for each level of eccentricity by dividing it by the highest eccentricity-related layer response. The normalization step was included to create an interpretable visualization of the eccentricity-related layer responses, as we were only interested in the relative differences per level of eccentricity, rather than the absolute layer response. To determine the preferred spatial frequency at each eccentricity level, we considered the spatial frequency that yielded the highest normalized response. We show a schematic overview of the method in Fig. 3a. Finally, we fitted an exponential decay function to the eccentricity-related spatial frequency preference data in 3c.

#### 4.3.3 Radial Bias

To investigate the presence of radial bias in the first convolutional layer, we exposed the network to sinusoidal gratings of eight orientations at a spatial frequency of 4 c/deg, with phase-offsets between −Π and Π in 32 equidistant steps. For each grating, V1 feature maps were averaged into a population map. Subsequent, all population maps belonging to stimuli with the same orientation were averaged, resulting in *orientation maps*. A schematic overview of the method can be seen in Fig. 4a. Using a polar coordinate system, each orientation map was divided into 16 equally sized bins, each covering 45° of polar angle with 22.5° overlap. To minimize confounding effects arising from stimulus edges [48] and vignetting [49], we excluded the outer 1.2° of eccentricity of the orientation maps before analyzing the layer activity. Per stimulus orientation, the mean activity of each bin was rescaled by dividing the bin values by the highest bin activity, and taken as the averaged directional cell activity. The orientation bias for a given bar orientation is indicated by the bin with the highest value. A schematic overview of the method is shown in Fig.4.

We aimed to measure the presence and direction of an orientation bias, indicating a preference for a specific orientation in the V1 layer, by computing the ratio of the total unit activity along the grating orientation axis to the total unit activity along the orthogonal axis. We calculated the ratio *A_grating_/A_orthogonal_*by dividing the sum of average unit activities in the bins with a 0°and 180°angle to the grating orientation, denoted by *A_grating_*, by the sum of average unit activities in the bins with a 90°and 270°angle to the grating orientation, denoted by *A_orthogonal_*. If the ratio is close to one, it indicates no orientation bias. If it is greater than one, it indicates a radial bias, and if it is smaller than one, it indicates an orthogonal bias. The radial bias ratios for various grating orientations at a spatial frequency of 4 c/deg were computed and presented in Fig. 4c.

We repeated the original radial bias experiment while varying the spatial frequency content of the gratings (0.10, 0.25, 0.50, 1.00, 2.00, 3.00, 4.00, 5.00, 6.00 c/deg), and computed the average radial bias ratio per level of spatial frequency. The radial bias per level of spatial frequency is computed by first calculating the radial bias ratio per stimulus orientation. We then average the radial bias ratio across orientations per level of spatial frequency to obtain the radial bias ratio per level of spatial frequency. The results are displayed in Fig.5.

When moving in a orthogonal direction from the radial axis of the stimuli, stimulus edges become more curved due to the RGC sampling-related distortion. We investigated whether this eccentricity-related progressive curving introduced by the RGC sampling resulted in sub-optimal V1 filter responses. To do this, we first created seven-by-seven-by-three achromatic lines varying in curvature. The straight lines were white on a black background, had a width of one pixel, and varied in orientation (0°, 45°, 90°, and 135°). We then introduced curving to the lines six steps, going from straight to maximum curvature given the amount of available pixels. Curving was introduced in both directions orthogonal to the line’s orientation. In addition, we created 3D Gabor kernels with the wavelength of the sinusoidal factor set to 2*π*, and with orientations from 0° to 180° degrees, in 16 steps. Next, we measured the cosine similarity of the Gabor and RSL-CORnet-Z filters with the curvature stimuli, per level of line curvature. Results are displayed in Fig.6a.

We investigated the role of RGC-sampling in generating the observed orientation bias in the first convolutional layer. The RSL layer, which performs the RGC-sampling, introduces a non-uniform distortion of the visual input resulting in eccentricity-related changes in spatial frequency. Additionally, edges in sinusoidal gratings are compressed in non-radial directions. To explore whether these local variations in spatial frequency could account for the orientation bias in V1, we determined the spatial frequency sensitivity of the layer. To accomplish this, we presented achromatic seven-by-seven-by-three sinusoidal gratings varying in spatial frequency (0.10 to 3.00 cycles per image), orientation(zero to *π* in 8 steps), and phase-offset (zero to 2*π* in 12 steps). The gratings were processed by each V1 filter, followed by the ReLU activation function, and the maximum activity for all gratings was averaged per level of spatial frequency. Our results, presented in Fig. 6b, demonstrate the spatial frequency sensitivity of the V1 layer.

To analyze the local spatial frequency content of grating stimuli in retinal space, we employed Gabor wavelet analysis. This technique involved generating a set of Gabor wavelets with varying frequencies and orientations. We then convolved each (resampled) grating stimulus with every wavelet in the set. By examining the response of each wavelet at every pixel of the stimulus, we assigned a dominant frequency to each pixel based on the wavelet that elicited the largest response. This allowed us to obtain a detailed map of the local frequency content of the grating stimuli in retinal space. We combined the local frequency content of the grating with the spatial frequency sensitivity of the V1 layer to determine the local layer response to the gratings. Two gratings with 45°orientation, and 0.1 and 5.0 c/deg respectively were tested (Fig. 6c). We again used the polar bin method on the resulting estimated V1 layer responses (Fig. 6d, right column). The results are displayed in Fig. 6e. Additionally, min-max normalized results are displayed in Fig. 6f.

## Supplementary information

**Supplementary Figure 1.**
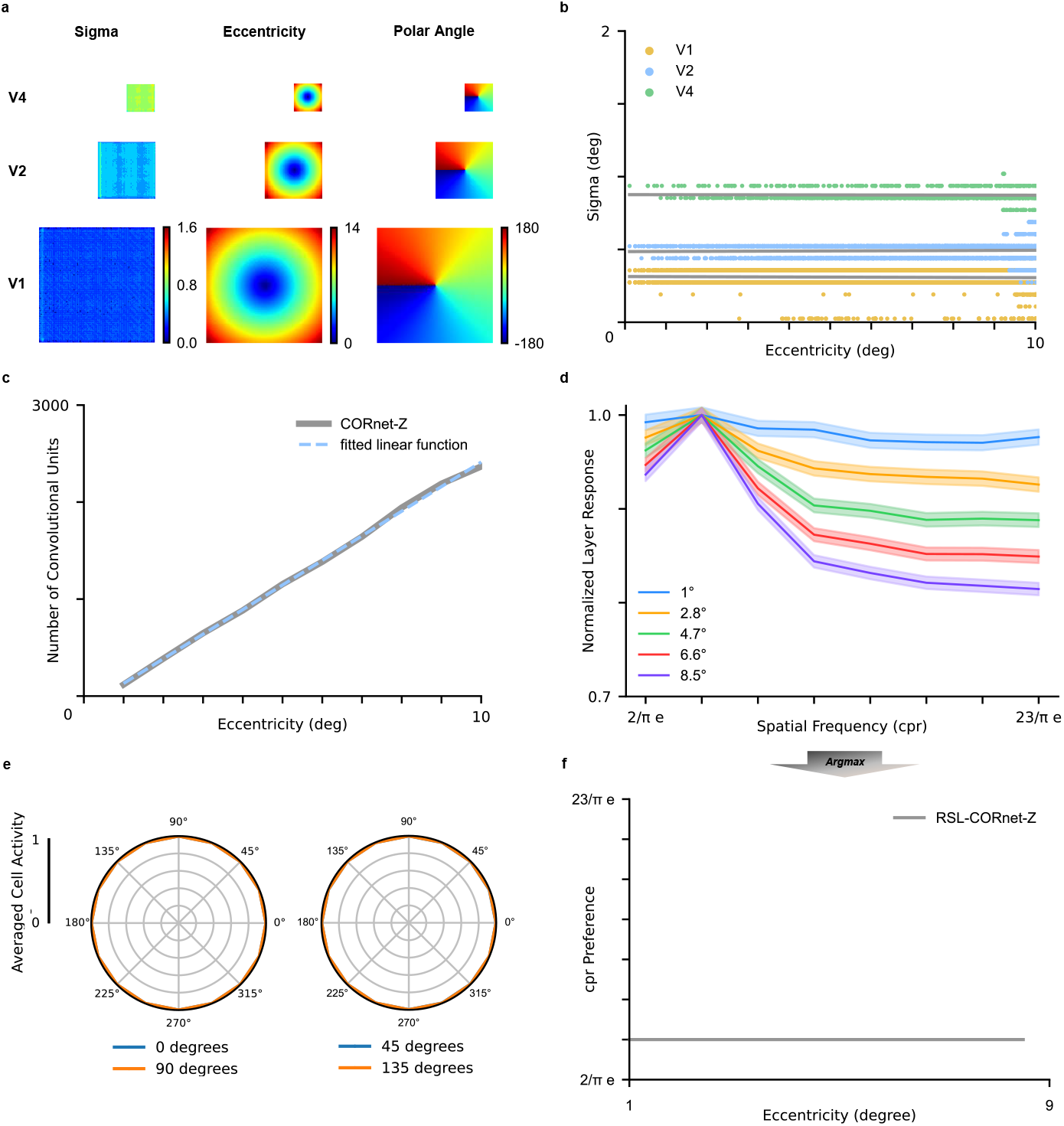
Results for CORnet-Z. **a**, Retinotopic maps of convolutional layers. **b**, Receptive field size as a function of eccentricity for V1, V2, and V4 layers. **c**, Convolutional units per level of eccentricity for the V1 layer. **d**, Normalized averaged layer response for the V1 layer as a function of spatial frequency. **e**, Polar plots displaying radial bias for 4 c/deg sinusoidal gratings. The left plot shows the result for 0° vs 90° sinusoidal gratings, the right plot shows 45° vs 135°. **f** Spatial frequency preference as a function of eccentricity.

**Supplementary Figure 2.**
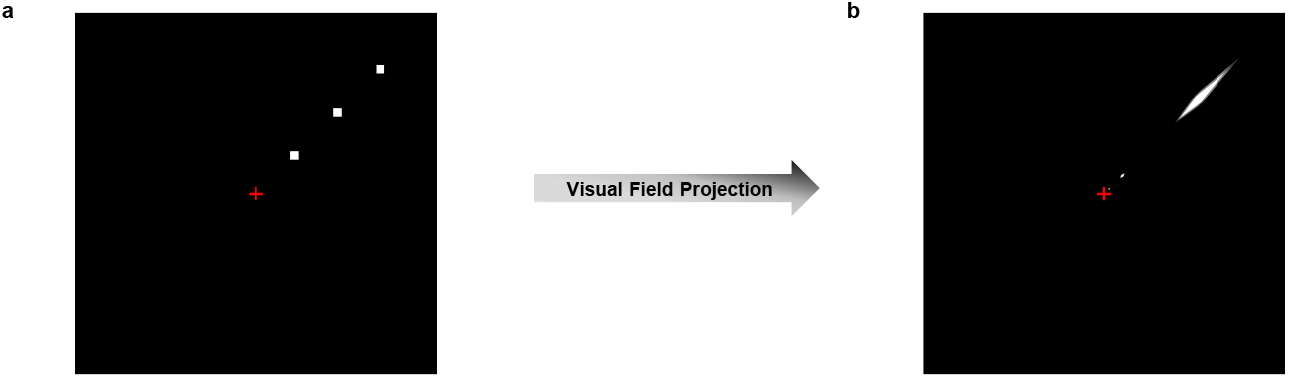
Visualization of receptive fields. **a**, Visualization of V1 receptive fields. Sampling is performed on the output of the RSL layer. **b**, Identical receptive fields as in a, but projected in the visual field. Receptive fields are elongated towards the fovea. The amount of elongation is a function of eccentricity.

**Supplementary Figure 3.**
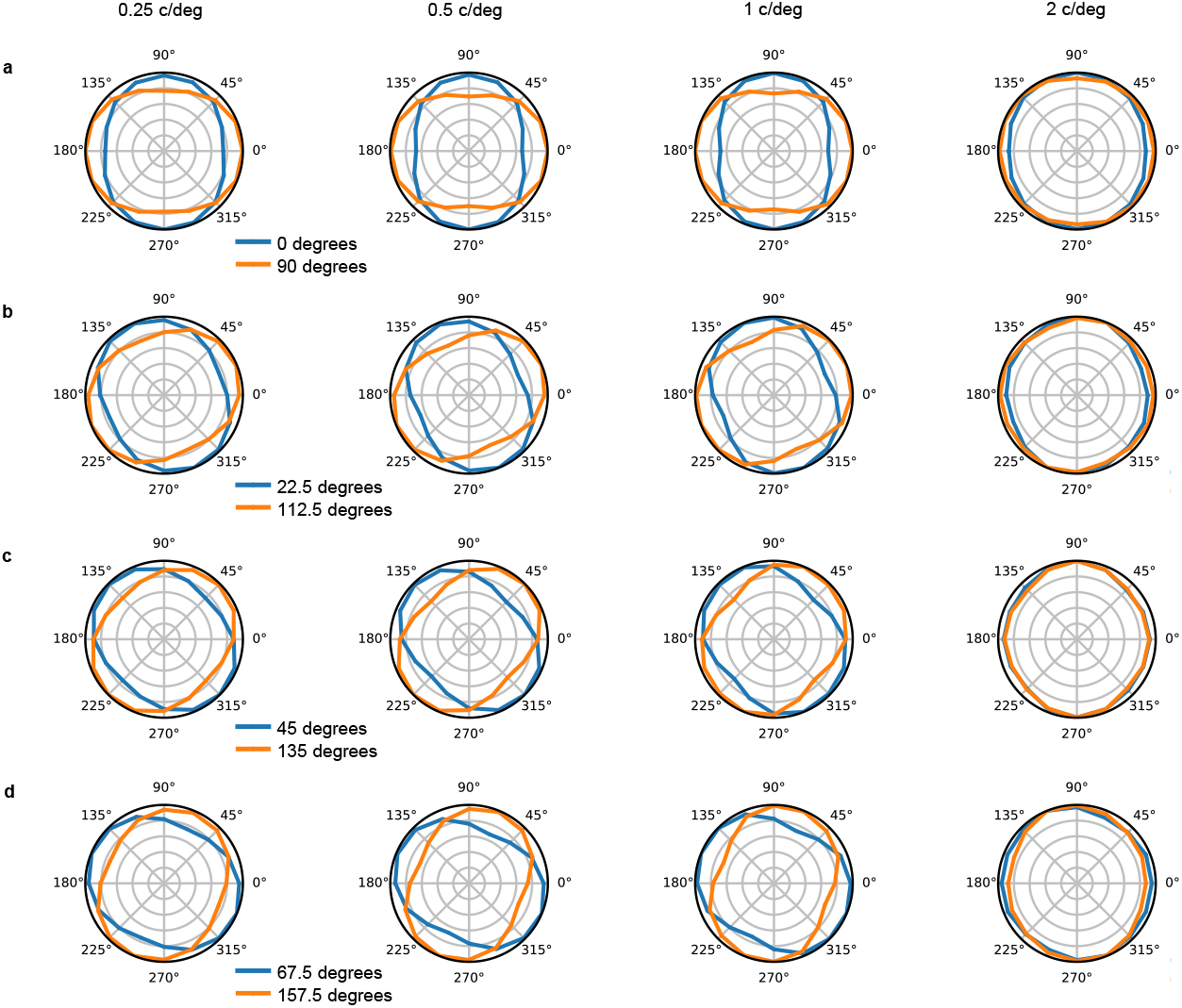
RSL-CORnet-Z radial bias polar plots for lower spatial frequencies. The columns show the results for 0.25, 0.50, 1, and 2 c/deg gratings, respectively. **a**, Polar plots show the results for 0° vs 90° sinusoidal gratings. **b**, Results for 22.50° vs 112.50° gratings. **c**, Polar plots for 45° vs 135° gratings. **d**, Results for 67.5° vs 157.5° gratings.

**Supplementary Figure 4.**
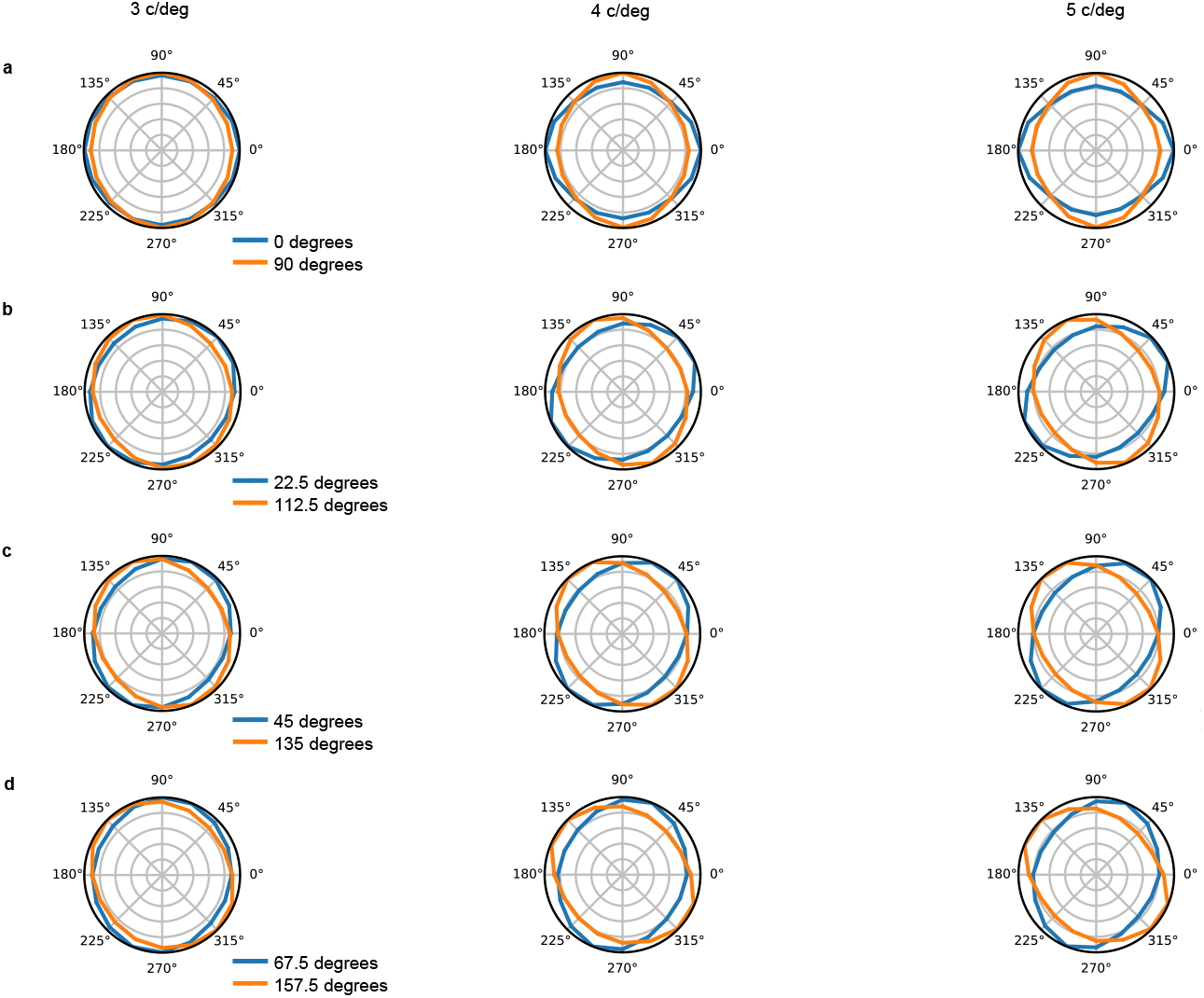
RSL-CORnet-Z radial bias polar plots for the higher spatial frequencies. The columns show the results for 3, 4, and 5 c/deg gratings, respectively. **a**, Polar plots show the results for 0° vs 90° sinusoidal gratings. **b**, Results for 22.50° vs 112.50° gratings. **c**, Polar plots for 45° vs 135° gratings. **d**, Results for 67.5° vs 157.5° gratings.

**Supplementary Figure 5.**
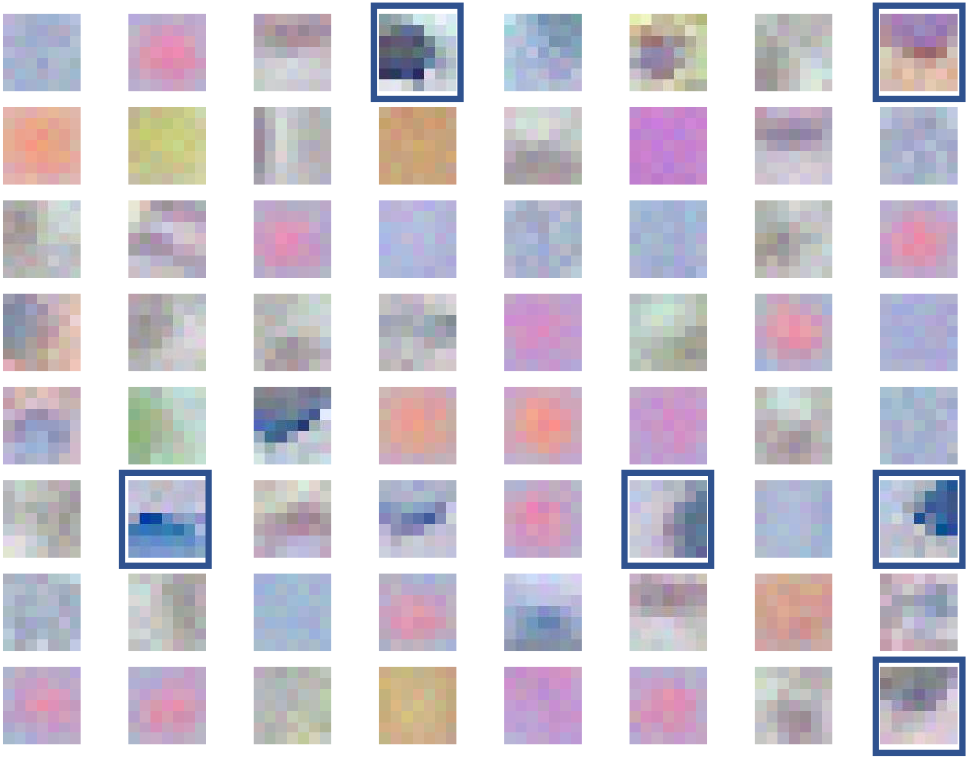
Visualization of V1 filters for RSL-CORnet-Z. V1 convolutional filters exhibit a curved profile to compensate for non-uniform distortion in visual input. Some filters show a notable curvature, which is highlighted by blue boxes in the image. Note that the extent of filter curvature is subjective.

## Acknowledgments

This study was funded by the European Union’s Horizon 2020 Framework under the Specific Grant Agreement Nos. 785907 (HBP SGA2) and 945539 (HBP SGA3).

